# Full cell infiltration and thick tissue formation *in vivo* in tailored electrospun scaffolds

**DOI:** 10.1101/2020.02.19.955948

**Authors:** Jip Zonderland, Silvia Rezzola, David Gomes, Sandra Camarero Espinosa, Ana Henriques Ferreira Lourenço, Andrada Serafim, Izabela Cristina Stancu, David Koper, Hong Liu, Pamela Habibovic, Peter Kessler, Marloes Peters, Peter Emans, Nicole Bouvy, Paul Wieringa, Lorenzo Moroni

## Abstract

Electrospun (ESP) scaffolds are a promising type of tissue engineering constructs for large defects with limited depth. To form new functional tissue, the scaffolds need to be infiltrated with cells, which will deposit extracellular matrix. However, due to dense fiber packing and small pores, cell and tissue infiltration of ESP scaffolds is limited. Here, we combine two established methods, increasing fiber diameter and co-spinning sacrificial fibers, to create a porous ESP scaffold that allows robust tissue infiltration. Full cell infiltration across 2 mm thick scaffolds is seen 3 weeks after subcutaneous implantation in rats. After 6 weeks, the ESP scaffolds are almost fully filled with *de novo* tissue. Cell infiltration and tissue formation *in vivo* in this thickness has not been previously achieved. In addition, we propose a novel method for *in vitro* cell seeding to improve cell infiltration and a model to study 3D migration through a fibrous mesh. This easy approach to facilitate cell infiltration further improves previous efforts and could greatly aid tissue engineering approaches utilizing ESP scaffolds.

**Statement of significance:** Electrospinning creates highly porous scaffolds with nano- to micrometer sized fibers and are a promising candidate for a variety of tissue engineering applications. However, smaller fibers also create small pores which are difficult for cells to penetrate, restricting cells to the top layers of the scaffolds. Here, we have improved the cell infiltration by optimizing fiber diameter and by co-spinning a sacrificial polymer. We developed novel culture technique that can be used to improve cell seeding and to study cytokine driven 3D migration through fibrous meshes. After subcutaneous implantation, infiltration of tissue and cells was observed up to throughout up to 2 mm thick scaffolds. This depth of infiltration *in vivo* had not yet been reported for electrospun scaffolds. The scaffolds we present here can be used for *in vitro* studies of migration, and for tissue engineering in defects with a large surface area and limited depth.

## 1. Introduction

Electrospun (ESP) scaffolds are highly porous and consist of nano- or micrometer sized fibers of natural or synthetic polymers, mimicking the fibrous composition of tissue extra cellular matrix (ECM)^[1–3]^. ESP scaffolds provide more mechanical support than hydrogels and are more flexible than scaffolds produced by additive manufacturing, making them interesting for tissue engineering approaches^[4]^. Large ESP mats are easily produced but are often limited to a thickness of several mm due to delamination and charge distribution. This makes ESP scaffolds particularly interesting for defects with a large surface area, but limited depth. This includes skin patches^[5]^, corneal repair^[6]^, cartilage regeneration^[7]^, vascular grafts^[8]^ and nerve guides^[9]^, among others. However, due to dense fiber packing and small pores, deep cell infiltration in ESP scaffolds remains a challenge^[10, 11]^. To create new fully functional tissue, ESP scaffolds first need to be fully infiltrated with cells. Several approaches have been developed to increase the cellular infiltration of ESP scaffolds, including increasing fiber diameter ^[12, 13]^, incorporating sacrificial salt-^[14]^ or ice crystals ^[15, 16]^ or co-spinning sacrificial polymer particles or fibers^[17–20]^. However, tissue infiltration *in vivo* has been limited to approximately 1 mm scaffold thickness. Increasing infiltration in a reliable and reproducible manner in scaffolds thicker than 1 mm remains a challenge^[17, 19]^.

Here, we combined two approaches to improve scaffold infiltration, by increasing fiber diameter and co-spinning sacrificial poly(ethylene glycol) (PEG) fibers. Using this method, we optimized porosity *in vitro* and achieved cell and tissue infiltration in up to 2 mm thick scaffolds *in vivo*. In addition, we propose a novel method to improve *in vitro* ESP scaffold cell seeding, and a novel 3D migration platform. Traditional methods to investigate migration are mostly limited to 2D substrates or 3D hydrogels^[21]^. We developed a transwell system that can guide cell migration to improve cell loading of ESP scaffolds, and could be used to research cytokine-driven 3D cell migration through a fibrous mesh.

## 2. Methods

### 2.1. Scaffold production

Electrospun (ESP) scaffolds were produced using the 300PEOT55PBT45 (PolyVation) polymer. 300PEOT55PBT45 was made by PolyVation from a starting 300 kDa poly(ethylene glycol) in the synthesis reaction, with a PEOT/PBT weight ratio of 55/45. A 20, 30 or 35% (w/v) 300PEOT55PBT45 solution was prepared in a mixture of 30% (v/v) 1,1,1,3,3,3-Hexafluoro-2-propanol AR (HFIP) (Bio-Solve) and 70% (v/v) Chloroform (Sigma-Aldrich) and dissolved under agitation overnight at room temperature. The PEG solution was 1.5% poly(ethylene oxide) (PEO) (Mw: 900,000 Da, Sigma-Aldrich) and 50% PEG (Mw: 3350 Da, Sigma-Aldrich) in a mixture of 25% (v/v) miliQ water and 75% (v/v) methanol (Sigma-Aldrich). Unless stated otherwise, the processing parameters for 300PEOT55PBT45 were: 3 ml/h flow rate, 15 cm working distance, 40% humidity and 23-25° C. The needle of both the 300PEOT55PBT45 and PEG were charged between 10-25 kV. The collector was charged between −1 and −10 kV. For the PEG solution, the flowrate was 3 ml/h and 25 cm working distance. For *in vitro* analysis, ESP scaffolds were produced on a 19 cm diameter mandrel at 100 RPM rotation on a polyester mesh (FinishMat 6691 LL (40 g/m^2^), generously provided by Lantor B.V.) with 8 mm diameter circular holes, on top of aluminum foil. After electrospinning, the aluminum foil was removed and circular ESP scaffolds were punched out with a diameter of 12 mm. Using this method, 12 mm ESP scaffolds were produced with a surrounding 1.5 mm polyester support ring to improve handleability. Unless stated otherwise, the ESP scaffolds used for *in vitro* analysis were 50μm thick. Different thicknesses were prepared by increasing the spinning time. Thickness was analyzed by cutting scaffolds in liquid nitrogen and analyzing the cross-section with SEM. The scaffolds for *in vivo* implantation were 300 μm thick and were produced on aluminum foil, without the polyester mesh. The aluminum foil was removed and discs were punched out with a 8 mm diameter. To dissolve the PEG solution, the scaffolds (and the 300PEOT55PBT45 only scaffolds to which they were compared) were incubated overnight in miliQ water at 50 °C. The next day, scaffolds were washed 5 times with water.

For sterilization for *in vitro* experiments, ESP scaffolds were submerged in 70% ethanol for 15 min and subsequently dried until visually dry. For sterilization for *in vivo* experiments, scaffolds were submitted to 254 nm UV light for 2 hours in vacuum.

### 2.2. Pore and fiber size quantification

Fiber size and pore area were manually measured using a custom-built Fiji script. 10-20 fibers or pores were selected in 5 different images of at least 2 different scaffolds. For pore area analysis, high contrast images were taken to create a dark background of pores deeper than a few fiber layers. The pore area of these pores on the surface of the scaffolds were measured in the biggest pores in each image.

### 2.3. Nano-CT

Nano-CT scans were recorded on a SkyScan 2211 high-resolution X-Ray nanotomograph (Bruker MicroCT, Belgium). The membranes were fixed on the sample holder with dental clay. All samples were scanned in nanofocus mode, using a CCD camera with a resolution of 4032 x 2688 and a pixel size of 0.5 μm. The source voltage and current were set at 30kV and 450 μA, respectively. The images were registered with a rotation step of 0.1 ° and an averaging of 4 frames at an exposure time of 1300 ms, resulting in a scanning time of approximately 5 hours for each sample. The resulted cross-sections were processed using CT NRecon software and subsequently reconstructed using CTVox. DataViewer software was used for analyzing the projections of the samples. CTAn software was used in order to obtain quantitative data regarding the porosity and wall thickness distribution of the analyzed samples. The analysis was performed in triplicate, on equal volumes of interest (VOI). All scanning, reconstruction, visualization and analysis parameters were kept constant for the analyzed samples.

### 2.4. Mechanical tests

The traction tests were performed using a Brookfield CT3 texture analyzer equipped with a 4500 g cell and a dual grip assembly (TA-DGA). The samples were cut at approximately 60 x 15 mm and tested at a speed of 0.5 mm/s, at room temperature. All measurements were performed in triplicate. A stress versus strain graph was plotted using the dedicated software and Young’s modulus was computed from the slope of the linear part of the traction curve, at 2% strain.

### 2.5. Cell culture

Bone marrow was isolated from a 22-year old male by aspiration by Texas A&M Health Science Center after ethical approval from the local and national authorities and written consent from the donor. Mononuclear cells were separated by centrifugation. Human mesenchymal stem cells (hMSCs) were subsequently isolated as described previously ^[22]^. Isolated hMSCs were received at passage 1 and tested for trilineage differentiation capacity. hMSCs were further expanded by seeding at 1000 cells/cm^2^ in tissue culture flasks in αMEM+Glutamax medium (Thermo Fisher Scientific) supplemented with 10% (V/V) fetal bovine serum (FBS) (Sigma-Aldrich) (basic medium) at 37 °C in 5% CO_2_. Cells were passaged at 70-80% confluency using 0.05% Trypsin and 0.53 mM EDTA (Thermo Fisher Scientific) and seeded on the ESP scaffolds at passage 5.

### 2.6. Cell migration quantification in vitro

To analyze cell infiltration, ESP scaffolds were placed in the bottom of a 48 well, or a 12mm transwell with 3μm pores in a polyester membrane insert (Corning). Rubber O-rings (outer diameter 12 mm, inner diameter 8 mm, Eriks) were placed on top of the scaffolds to prevent cells from reaching the bottom in any other way than through the ESP scaffolds. 15,000 hMSCs were then seeded on top of the scaffolds in basic medium and cultured for 4 days, unless stated otherwise. For the samples on transwells, 24 hours after seeding, medium in the top compartment was changed to medium without FBS and basic medium in the bottom. The medium was changed every 24 hours to sustain an FBS gradient. Cells were fixed with 3.6% (v/v) paraformaldehyde (Sigma-Aldrich) in PBS at room temperature for 20 minutes. DAPI (Sigma-Aldrich, 0.14 μg/ml in PBS+0.05% (v/v) tween-20) was then used to visualize the cells. Quantification of cells was done in 5 separate images of each of 3 different scaffolds.

### 2.7. In vivo subcutaneous implantation

All experiments and protocols were approved by the Dutch Central Committee for Animal Experiments (in Dutch: Centrale Commissie Dierproeven). Female rats were obtained (Crl:NIH-Foxn1rnu, 8-10 weeks old, 140-212g) (Charles-River) and housed at 21 °C with a 12 h light/dark cycle and had ad libitum access to water and food. Prior to anesthesia, buprenorphine 0.05 mg/kg bodyweight and carprofen 4 mg/kg were administered as premedication. The animals were subsequently anesthetized with isoflurane 3-4% (v/v) for induction, and isoflurane 2% (v/v) for maintenance, adjusted according to the clinical signs during surgery. After shaving, disinfection and draping of the animal’s dorsum, four 1 cm long linear skin incisions parallel to the spine were made, two on each side. Four subcutaneous pockets of maximum 10 mm x 10 mm were created. ESP scaffolds, 8mm diameter, that had been seeded on both sides of the scaffold 24h prior to implantation with a total of 15,000 hMSCs, were carefully placed inside the pockets. Scaffolds were randomly assigned to a pocket. After implantation, the skin was closed intracutaneously with Monocryl 4×0 sutures (Ethicon). Buprenorphine 0.03 mg/kg bodyweight was administered 8 hours after surgery. The morning of post-operative day 1 and 2, each animal was administered a dose of 4 mg/kg bodyweight carprofen. Thereafter, animal welfare was evaluated on a daily basis with a discomfort logbook scoring system, and appropriate medication was given only when needed. After 3- and 6 weeks, the animals were euthanized with gradual CO_2_ overdose. The sample and surrounding tissues were collected and processed for histology. No animal was lost during this study.

### 2.8. Tissue preparation and infiltration quantification

3 or 6 weeks post-implantation, skin samples containing the scaffolds were explanted. Tissue explants were cut with surgical scissors to fit the dimensions of the silicon molds (2×2×2 cm), without disrupting the generated pocket that contained the implants. Samples were then placed in 50 mL centrifugation tubes and fixed for 24h at 4 °C in a 3.6% (v/v) solution of paraformaldehyde in TBS (tris-buffered saline). After fixation, the samples were transferred to new tubes and underwent embedding in a series of: 30% (w/v) sucrose, then 50:50 volume ratio of 30 % sucrose and optimal cutting temperature compound (OCT) (Thermo Fisher Scientific), and lastly OCT only for 24h each. The samples were maintained in OCT until freezing. Tissue explant were placed inside silicon molds and the molds were filled with OCT. Freezing was conducted on the liquid-vapor interface of a liquid nitrogen tank to avoid formation of bubbles. Cross-sections of 7 μm thick were cut on a cryotome and samples were stained with hematoxylin and eosin (H&E). Sections were hydrated in deionized water and placed in Gill’s hematolxylin (III) (Sigma-Aldrich) for 5 min, in running water for 5 min, dehydrated and counterstained with alcoholic Eosin Y (Sigma-Aldrich) for 1 min. Sections were differentiated in 100% ethanol, allowed to air dry and mounted in DPX (Sigma-Aldrich).

The percentage of tissue infiltration was quantified by measuring the area of infiltrated tissue in the scaffold, divided by the total area of the scaffold. The infiltrated area was defined as clear dark staining in the H&E staining of the sections, where cells were surrounded by ECM. Areas where cells were present, but individual cells could still be distinguished without ECM formation in between, were considered non-infiltrated. This was measured in sections of 6-8 different scaffolds per condition.

### 2.9. Statistical analysis

The number of replicates is stated in the figure subtexts. Quantification of fiber diameter was done on randomly selected fibers. Pore area quantification was done on the biggest pores in each image. Scaffolds from each condition were randomly assigned to a pocket for *in vivo* implantation. Normal distribution of each experimental group was tested with the

Shapiro-Wilk test. Statistical significance was tested with a One-way ANOVA with Tukey’s post hoc for experiments with multiple comparisons, or two-tailed student’s t-test for experiments with one comparison. Graphpad Prism 8 was used to perform statistical analysis and significance was set at p<0.05.

## 3. Results

### 3.1. Optimizing scaffold porosity by increasing fiber diameter

Increasing fiber diameter has been shown to increase ESP scaffold porosity and pore size^[12, 13]^. To increase the fiber diameter of the 300PEOT55PBT45 ESP scaffolds, we evaluated the effect of working distance, flow rate and polymer concentration. The fiber diameter increased most by increasing the polymer concentration (Fig 1a, b). Flow rate had a small effect on the fiber diameter (Fig. 1a), while the working distance had no effect (Supplementary Fig. 1). Fiber diameter increased from 1.1±0.3 μm with 20% at 3 ml/h to 3.3±0.4 with 35% at 3ml/h. These scaffolds were used for further analysis. Pore size on the surface of the scaffolds was analyzed and showed an increase from 8.3±2.3 μm^2^ in the 20% to 19.5±5.4 μm^2^ in the 35% scaffolds (Fig. 1c). Next, cell migration through 50μm thick scaffolds was analyzed. hMSCs were seeded on top of the scaffolds and were cultured for 1, 4 and 7 days. Cells were fixed, stained with DAPI and the number of cells on the bottom of the scaffold was quantified. On day 1, no cells were found at the bottom of the 20% or 35% scaffolds, showing that cells did not fall through the scaffold and could only reach the bottom by migration. On day 4 and 7, almost no cells (<1 cell/mm^2^) were found on the bottom of the 20% scaffolds. However, 47.1±6.4 and 50.7±12.8 cells/mm^2^ were found on day 4 and 7 on the bottom of the 35% scaffolds, respectively. This demonstrates that by increasing the fiber diameter, the pore space could be sufficiently increased to allow migration through the 50μm scaffolds. As no difference was found between day 4 and 7, and no cells were observed on day 1, all migration experiments after this were measured at day 4.

**Figure 1.**
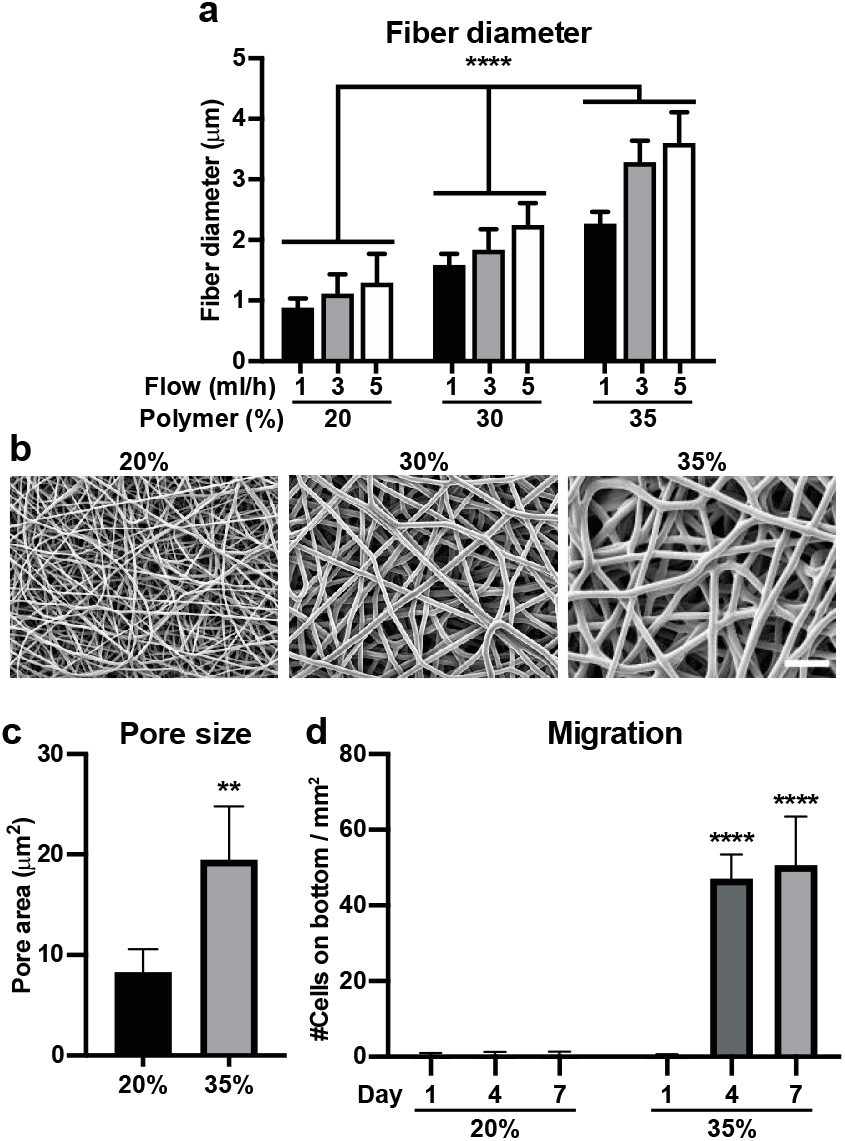
Optimizing fiber diameter and scaffold porosity for cell migration. **a,** Different concentrations of 300PEOT45PBT55 (w/v) were electrospun at different flow rates. Significance indicates differences between polymer concentrations at the same flow rate. **b,** SEM images of 20%, 30% and 35% (w/v) 300PEOT45PBT55 concentration spun at 3 ml/h. Scalebar 20μm. **c,** Estimation of pore size on the surface of the ESP scaffolds, analyzed from SEM images (average of 10-20 pores from 10 different images). **d,** Number of hMSCs on the bottom of 20% or 35% (w/v) 300PEOT45PBT55 scaffolds 1, 4 or 7 days after cell seeding on top of the scaffolds. Cells were counted on 5 different images of each of 3 scaffolds. Significance indicates statistical differences between 35% day 1, and the same day of 20%. **b, d,** One-way Anova and **c,** Student’s t-test. ** p<0.01, **** p<0.0001. Error bars indicate mean±SD.

### 3.2. Further increasing porosity by co-spinning sacrificial fibers

Scaffold porosity has previously been increased by co-spinning sacrificial electrospun fibers^[17–20]^. These fibers take up space during the electrospinning process and are later dissolved, leaving extra empty space in the scaffolds. To further improve the porosity of our scaffolds, we spun 20% and 35% 300PBT55PBT45 with one needle, and a PEG solution with a second needle. The PEG fibers were approximately 3 μm in diameter. After dissolving the PEG in water, surface pore area increased in both the 20% and 35% scaffolds (Fig. 2a, b). In the 20% scaffolds, pore size increased from 8.3±2.3 μm^2^ to 27.6±5.0 μm^2^ (p<0.0001), and from 19.5±5.4 μm^2^ to 43.5±10.2 μm^2^ (p<0.0001) in the 35% scaffolds. Mechanical properties were lower for the scaffolds created with PEG fibers (&PEG) (~2-3MPa), compared to the scaffolds created without PEG (~4-5MPa) (Supplementary Fig. 2a-c). This reduction in mechanical properties could be due to less inter-fiber linking in the scaffolds created with PEG fibers. Sacrificial fibers will interrupt the merging between fibers that typically occurs during electrospinning. So, the more porous mesh is less rigid, potentially more amendable to cell-mediated rearrangement and/or increased pore size due to swelling. We then analyzed the migration of hMSCs through 50μm thick scaffolds 4 days after seeding. Interestingly, for both the 20% and 35% scaffolds, no difference in cell infiltration was found between the scaffolds created with or without PEG. Similar cell numbers were found on the bottom of both 35% scaffolds (35% and 35% &PEG), while very few cells were found on the bottom of the both 20% scaffolds (20% and 20% &PEG).

**Figure 2.**
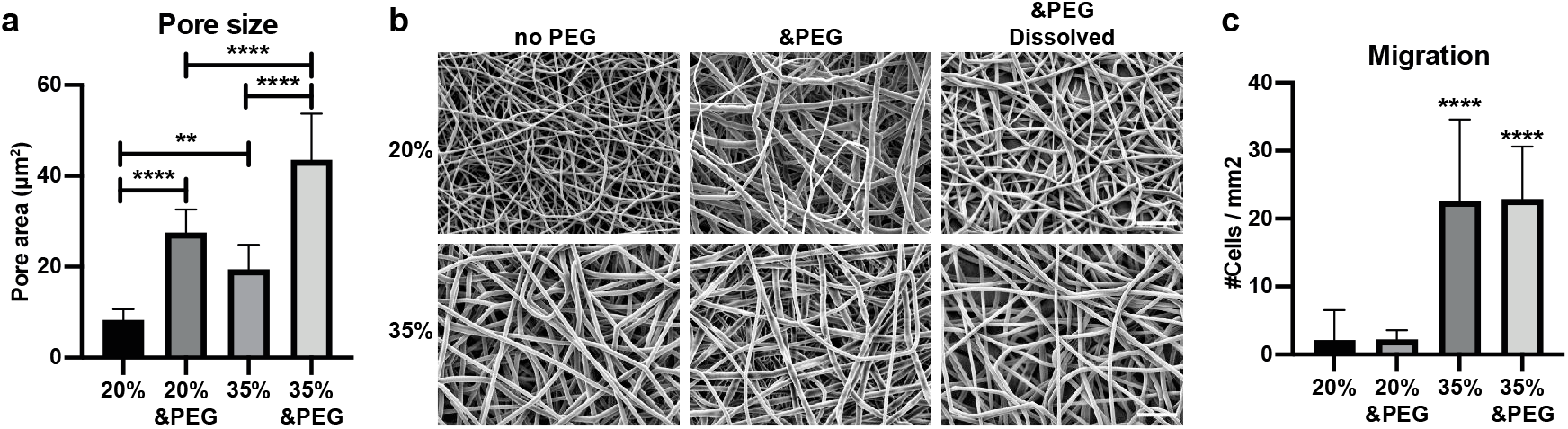
Improving scaffold porosity using a sacrificial polymer. **a,** Estimation of pore size on the surface, or **b,** SEM images, of ESP scaffolds created with 20% or 35% (w/v) 300PEOT45PEG55 with or without a second needle spinning a mixture of high and low Mw PEG. The PEG fibers were later dissolved in water. Analyzed from SEM images (average of 10-20 pores from 10 different images). Scalebar 20μm. **c,** Number of hMSCs on the bottom of the different scaffolds 4 days after cell seeding on top of the scaffolds. Cells were counted on 5 different images of each of 3 scaffolds. Stars indicate significant difference with both 20% and 20% &PEG. **a, c,** One-way Anova. ** p<0.05, **** p<0.0001. Error bars indicate mean±SD.

### 3.3. Guided migration reveals differences between scaffolds created with or without sacrificial fibers

To further investigate the differences between the 35% scaffolds created with or without sacrificial fibers and to investigate the limits of cell infiltration, we created scaffolds with thicknesses of 50, 100 and 150μm. Scaffolds were placed on the bottom of a normal cell culture well, as with the previous migration experiments, or on top of a transwell. In the normal cell culture well, cells migrated to the bottom of the 50μm scaffolds and no difference was found between the 35% and 35% &PEG scaffolds(Fig. 3a). No hMSCs migrated to the bottom of 100 μm or 150 μm thick scaffolds. In the transwell system, medium containing FBS was put in the bottom compartment and medium without FBS was put in the top compartment, creating a gradient to guide cell migration. Medium was refreshed every day to maintain the FBS gradient and the bottom of the scaffolds was analyzed after 4 days of migration. Interestingly, cells were found at the bottom of the 100 μm thick 35% scaffolds (Fig. 3b). Still, no cells migrated to the bottom of the 150 μm thick 35% scaffolds. In the 35% scaffolds created with PEG fibers, migration to the bottom of the scaffolds was increased for all thicknesses, compared to 35% scaffolds. Also, many cells were found on the bottom of the 150 μm thick scaffolds. These results show that the FBS gradient guided cell migration towards the bottom of the scaffold. This method revealed the differences in the scaffolds’ ability to allow infiltration. In addition, this novel method could be used to increase cell infiltration of ESP scaffolds, or as a tool to study cell migration through a fibrous mesh.

**Figure 3.**
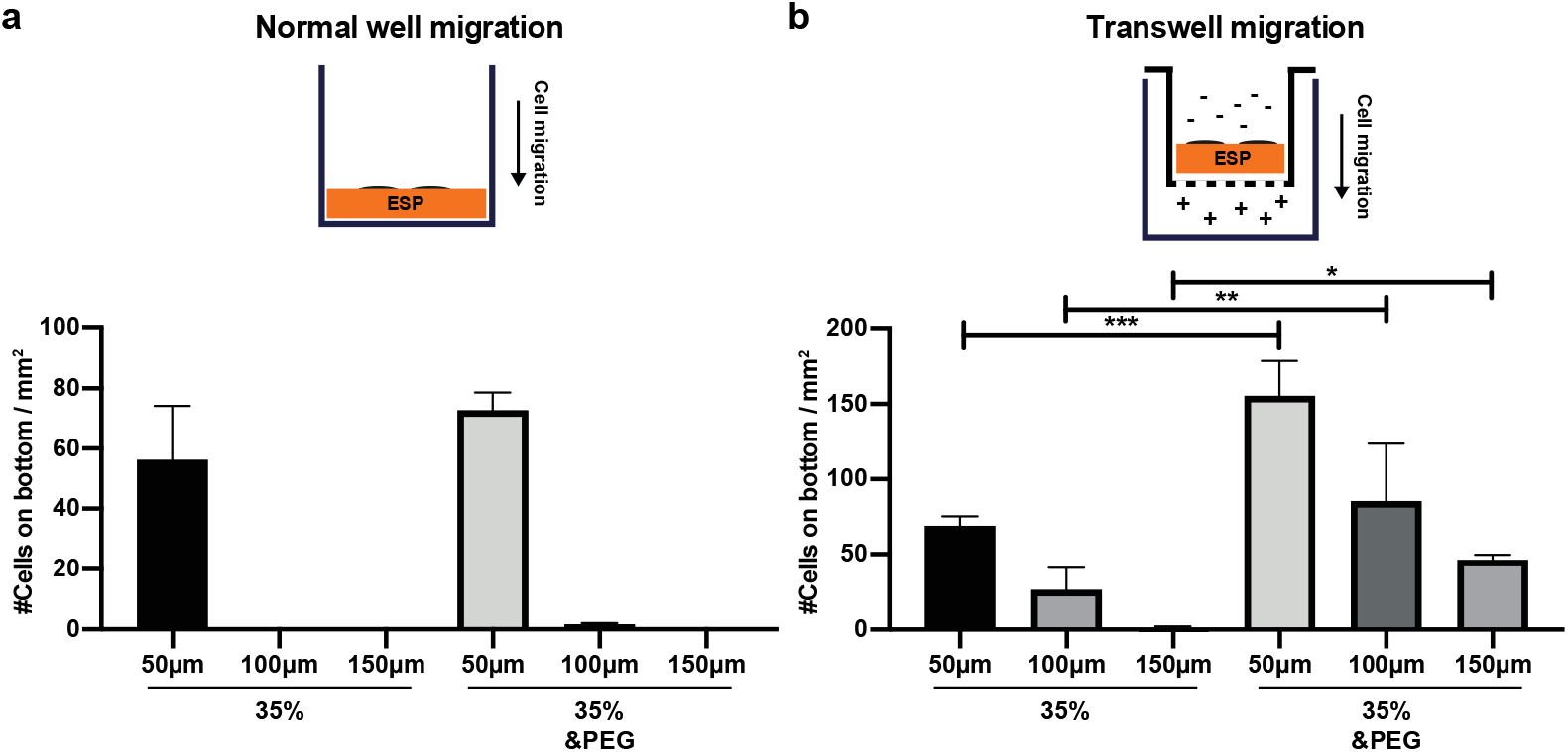
Migration through ESP scaffolds with and without cytokine attraction. **a, b** hMSCs on bottom of 50, 100 or 150μm thick ESP scaffolds, created with or without sacrificial PEG fibers, 4 days after seeding on top of the scaffolds. Cells were counted on 5 different images of each of 3 scaffolds. **a,** hMSCs were seeded on top of a ESP scaffold in the bottom of a normal 48-well cell culture well with medium containing FBS on top. **b,** hMSCs were seeded on top of an ESP scaffold placed inside a 3μm transwell system, with medium without FBS on top and medium with FBS in the bottom compartment. One-way Anova. * p<0.05, ** p<0.01, *** p<0.001, **** p<0.0001. Error bars indicate mean±SD.

### 3.4. Nano-CT analysis does not explain differences in scaffold infiltration

It is not fully understood what key aspects of an electrospun scaffold allow deep cell infiltration. To better describe our scaffolds, we analyzed the 20% and 35% scaffolds created with or without PEG fibers with nano-CT (Supplementary Fig. 3 and Supplementary Movie 1-4). Interestingly, the pore size distribution was different between the scaffolds with or without PEG, but not different between 20% and 35%, or between 20% &PEG and 35% &PEG (Fig. 4a). In the 20% and 35% scaffolds without PEG, most pores were in the 5μm diameter range, while both scaffolds with PEG had many pores with 10-15μm diameter. The volume and percentage of closed pores was significantly reduced in the 20% &PEG, 35% and 35% &PEG scaffolds, compared to the 20% scaffold (Fig. 4b, c). However, the percentage of closed pores was lower than 0.0005% of total volume in all scaffolds, so unlikely to greatly affect the cell migration. Lastly, the total pore volume and porosity was slightly higher in the 20% &PEG and 35% &PEG scaffolds, but all scaffolds had a porosity of around 70-80% (Fig. 4d, e). While a big difference in cell infiltration was found between the different scaffolds (Fig. 2c, Fig. 3a, b), no correlation was found with the nano-CT results. This highlights that porosity and pore size alone cannot explain the ability of an electrospun scaffold to allow cell infiltration.

**Figure 4.**
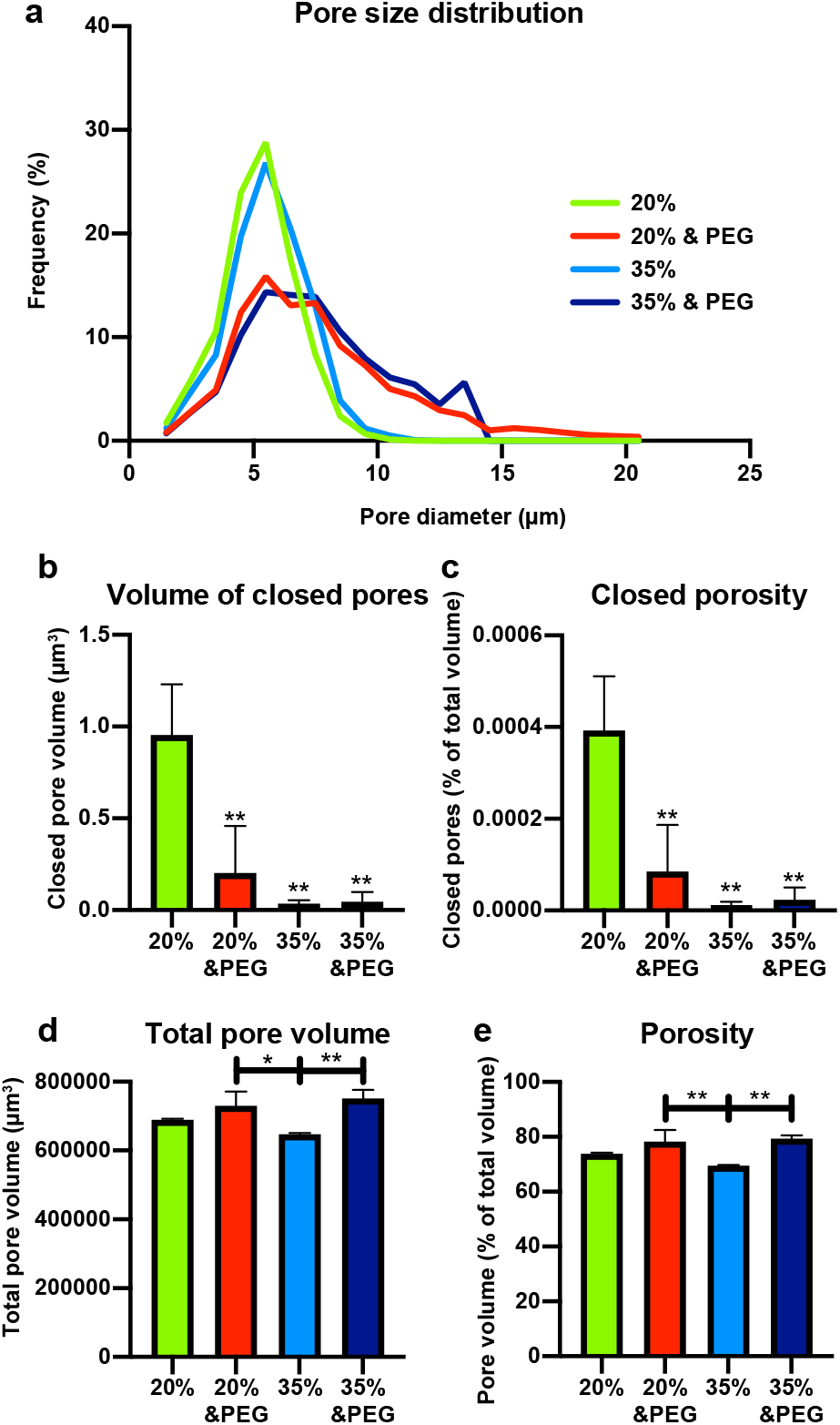
Nano-CT analysis of the ESP scaffolds. 20% and 35% 300PEOT45PBT55 scaffolds, created with or without sacrificial PEG fibers, were analyzed with nano-CT for **a,** pore size distribution, **b,** closed pore volume and **c,** closed porosity, **d,** total pore space and **e,** porosity. n=3 scaffolds for each condition.

### 3.5. Full cell infiltration and tissue formation in subcutaneously implanted scaffolds

To further test the ability of the ESP scaffolds to allow for cell infiltration, we implanted 300 μm thick 35% and 35% &PEG scaffolds in subcutaneous pockets of immunodeficient rats. We seeded the scaffolds with hMSCs and compared to cell-free scaffolds, to see the effect of the hMSCs on tissue infiltration. After 3 weeks, cells were present throughout the whole thickness of all scaffolds (Fig. 5a, b). The amount of tissue formation, however, was significantly increased in the 35% &PEG scaffolds, compared to the 35% scaffolds (Fig. 5c). Around 50% of the 35% &PEG scaffolds was filled with *de novo* tissue, compared to ~25% in the 35% scaffolds. No difference was found between hMSC-seeded scaffolds and cell-free scaffolds. After 6 weeks, the differences were even more pronounced, and tissue formation increased for both the 35% and 35% &PEG scaffolds, compared to 3 weeks. The 35% &PEG scaffolds were on average filled with >80% tissue, compared to the <50% of the 35% scaffolds. Again, no difference was found between cell-laden and cell-free scaffolds. Interestingly, the thickness of all scaffolds significantly increased after implantation. While 300 μm thick scaffolds were implanted, scaffolds were up to 2 mm thick after explantation (Supplementary Fig. 4). No significant differences in scaffold thickness were found between 3- and 6 weeks post-implantation, nor between different scaffold types, with or without cells. All groups averaged between 1.1 and 1.5 mm. This shows that these scaffolds allowed for great cell infiltration and tissue formation to up to 2 mm, something not previously reported for electrospun scaffolds.

**Figure 5.**
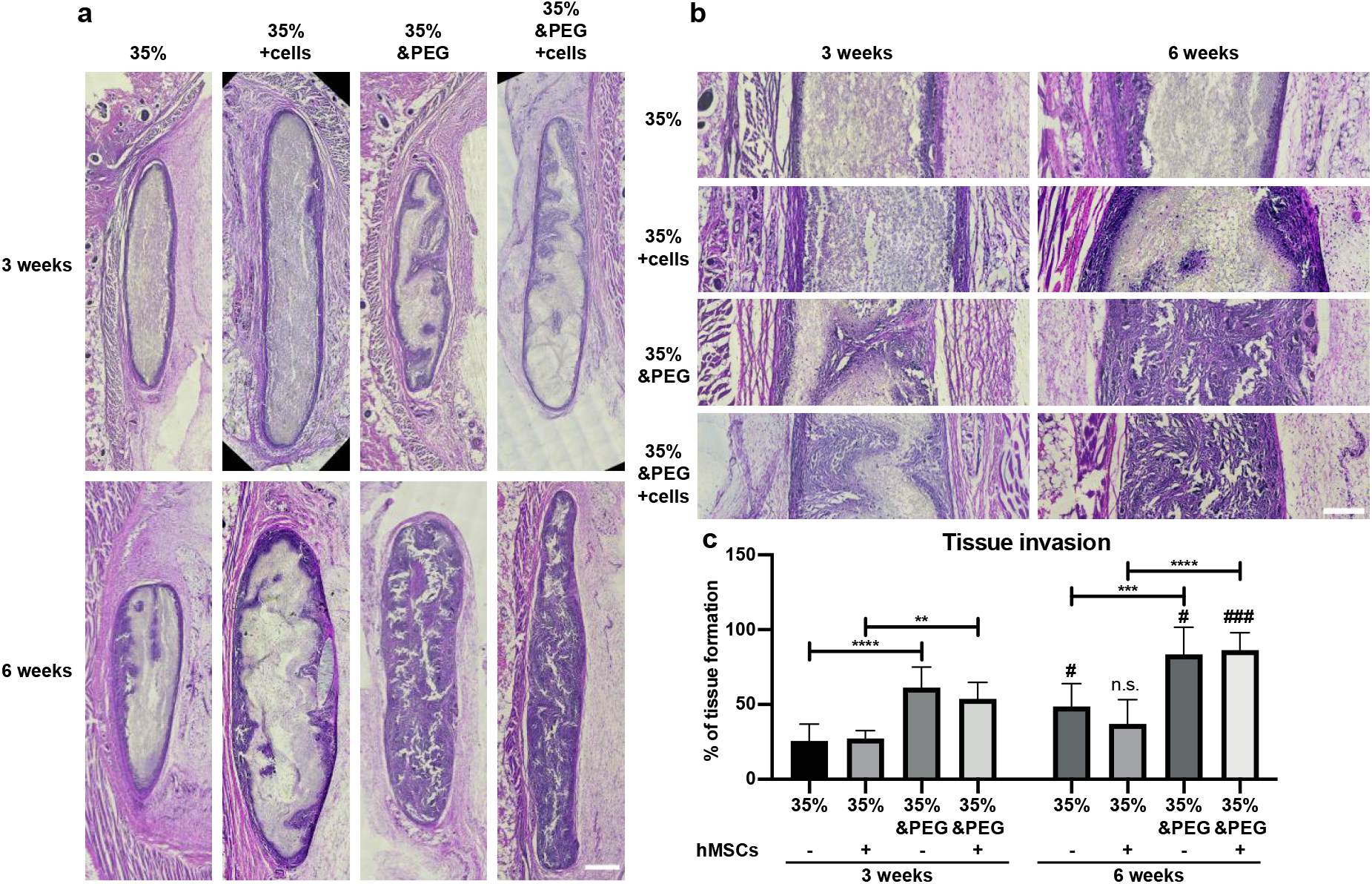
Cell infiltration and tissue formation of subcutaneously implanted ESP scaffolds. **a,** 35% 300PEOT45PBT55 scaffolds, created with or without sacrificial PEG fibers, were implanted in subcutaneous pockets and analyzed for infiltration after 3 (top panels) or 6 (bottom panels) weeks. Scaffolds were seeded with hMSCs 24h prior to implantation and compared to scaffolds without cells. Scalebar 500μm. **b,**Detail of the ESP scaffolds from the respective images in **a**. **c,** Quantification of tissue formation in the different scaffolds 3- and 6 weeks post-implantation. Asterisks indicate statistical difference between the indicated groups. Number signs indicate differences between 3- and 6 weeks of the same scaffold type. n=6-8 rats for each group. One-way Anova. ** p<0.01, *** p<0.001, **** p<0.0001. Error bars indicate mean±SD.

## 4. Discussion

Scaffold porosity was optimized here by combing two established methods, e.g. increasing fiber diameter and adding sacrificial polymer fibers^[12, 13, 17–20]^. The Nano-CT analysis revealed differences in overall porosity, closed porosity and pore size distribution, but did not correlate with the scaffolds’ performance on the cell migration tests. This showed that these parameters do not fully describe the essential properties of ESP scaffolds that allow cells to migrate through them. Other attributes likely describe this better, such as the minimal pore size of a series of interconnected pores, and the complexity of this maze. We have recently shown that hMSCs coming from electrospun scaffolds can migrate through 3 μm pores^[23]^. The minimal pore size that cells can migrate through in electrospun scaffolds could be in a similar range. However, even using nano-CT, these scaffold attributes are difficult to quantify. Novel algorithms and imaging techniques could improve this, greatly aiding ESP scaffold optimization and characterization.

Other reports have also shown robust ESP infiltration *in vitro* and *in vivo*. This includes the use of salt crystals^[14]^, ice crystals^[15, 16]^ PEG fibers^[18]^, and PEG microparticles^[20]^ and wet spinning^[24]^. Infiltration of 150 μm, as we’ve demonstrated here, has been achieved by the addition of salt crystals after 3 weeks of culture^[14]^. However, we report 150 μm infiltration already after 4 days. Other reports demonstrate limited cell infiltration of 50-100 μm^[15, 16, 18, 25]^. 150 μm infiltration *in vitro* using PEG fibers has not been previously reported, but using PEG microparticles, 400 μm infiltration of fibroblasts was achieved after 4 days of culture^[20]^. However, different cell types have different migration properties^[21]^ and hMSCs have not been previously shown to deeply infiltrate an ESP scaffold in only 4 days, as we have shown. ~250 μm infiltration of hMSCs after 3 weeks of culture has been reported^[26]^.

Dynamic seeding of ESP scaffolds has resulted in the filling of 2.5 mm scaffolds *in vitro*^[24]^. This cannot be considered cell infiltration but could be a useful tool to increase cell distribution in ESP scaffolds before seeding. The transwell method presented here can also be used to increase cell infiltration before implantation. In addition, this method could be used to study cell migration through fibrous meshes. Cell migration is often studied in 2D, or in 3D hydrogels^[21]^. However, natural ECM is a fibrous matrix with fiber diameters ranging from nano-to micrometer sized fibers^[1–3]^. ESP scaffolds could be an interesting tool to study 3D migration, more closely mimicking the natural ECM.

Several attempts have previously been made to improve scaffold infiltration *in vivo*. Many ESP scaffolds only allow limited infiltration of 200-400 μ m ^[15, 16]^. Using PEG fibers, others have shown infiltration of tissue up to 1 m m ^[17, 19]^. The greatest infiltration of tissue in ESP scaffolds reported in literature is of 1.3 mm thick scaffolds, using PEG microparticles^[20]^. Here, we report infiltration of ESP scaffolds of up to 2 mm thick *in vivo*. The increase in size from 300 μm thick scaffolds upon implantation to 1-2 mm thick after 3 weeks could be attributed to cell infiltration and tissue formation that expand the scaffold. Seeding cells in a thinner scaffold that will later expand could be beneficial, as cells can be distributed through a 300 μm thick scaffold more easily than a thicker scaffold. The expansion in size would still allow for a thick layer of tissue to form. Others have also reported a slight change in scaffold size after implantation^[19]^, but not as significant of an increase as we report here. Scaffold properties such as fiber stiffness and strength of inter-fiber connections could, among others, potentially influence this expansion in size. Further research into this phenomenon could improve the tissue engineering approaches utilizing ESP scaffolds.

Here, we show the optimization of ESP scaffold porosity using an increase in fiber diameter and sacrificial PEG fibers. We propose a novel *in vitro* method to research cytokine-attracted 3D migration through fibrous meshes. Also, the ESP scaffolds created here allowed for cell infiltration *in vivo* of up to 2 mm, a thickness that has not previously been reported for ESP scaffolds.

## Acknowledgements

We are grateful to the European Research Council starting grant “Cell Hybridge” for financial support under the Horizon2020 framework program (Grant #637308). Some of the materials that were used in this work were provided by the Texas A&M Health Science Center College of Medicine Institute for Regenerative Medicine at Scott & White through a grant from NCRR of the NIH (Grant #P40RR017447). The nano-CT analyses were possible due to European Regional Development Fund through Competitiveness Operational Program 2014-2020, Priority axis 1, ID P_36_611, MySMIS code 107066, INOVABIOMED. This research has been made possible with the support of the Dutch Research Council (NWO, Grant #16711) and the Dutch Province of Limburg (LINK project).

## Supplementary Figures

**Supplementary figure 1.**
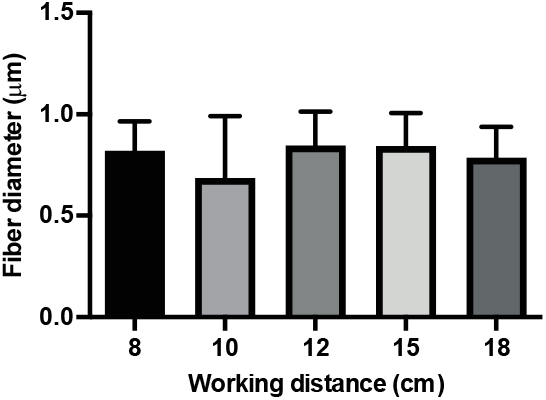
Effect of working distance on fiber diameter. 20% (w/v) 300PEOT45PBT55 was spun at 1 ml/h at different distances from the collector.

**Supplementary figure 2.**
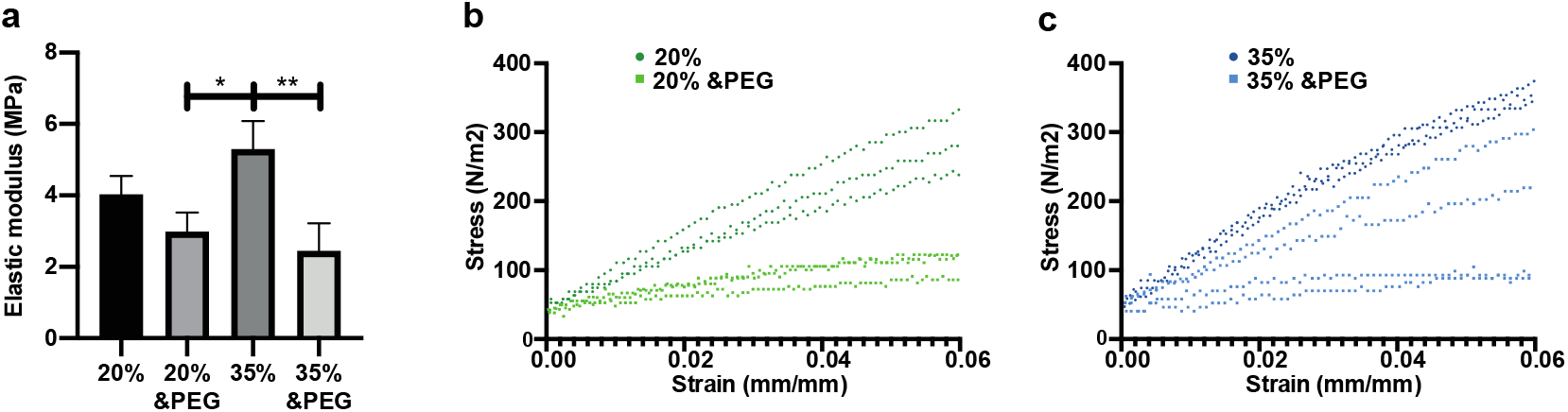
Mechanical properties of ESP scaffolds. **a,**Tensile mechanical tests of 20% and 35% 300PEOT55PBT45 scaffolds, created with or without addition sacrificial PEG fibers. One-way Anova. * p<0.05, ** p<0.01. **b,** Individual data points of each replica in the first 6% strain of 20% and 20% &PEG and **c,** 35% and 35% &PEG. n=3 for 20%, 20% &PEG and 35%, n=4 for 35% &PEG.

**Supplementary figure 3.**
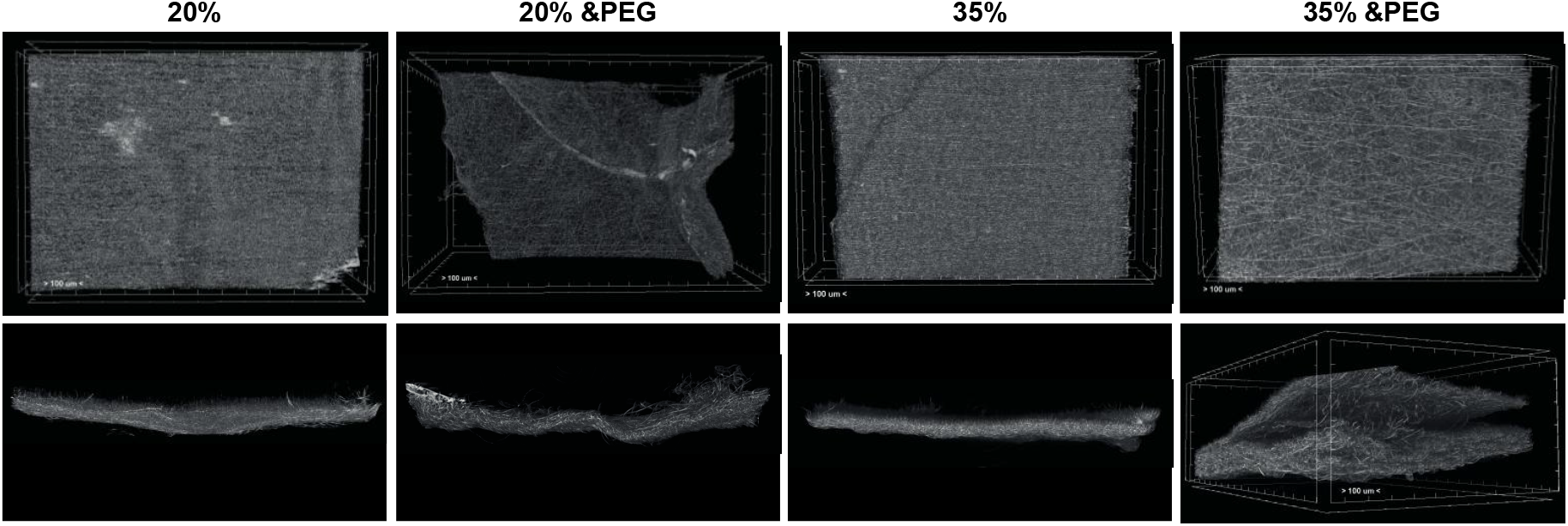
Nano-CT images of the different scaffolds. Top view in the top panels and side view in the bottom panels. Delaminated areas were excluded from the measurements.

**Supplementary figure 4.**
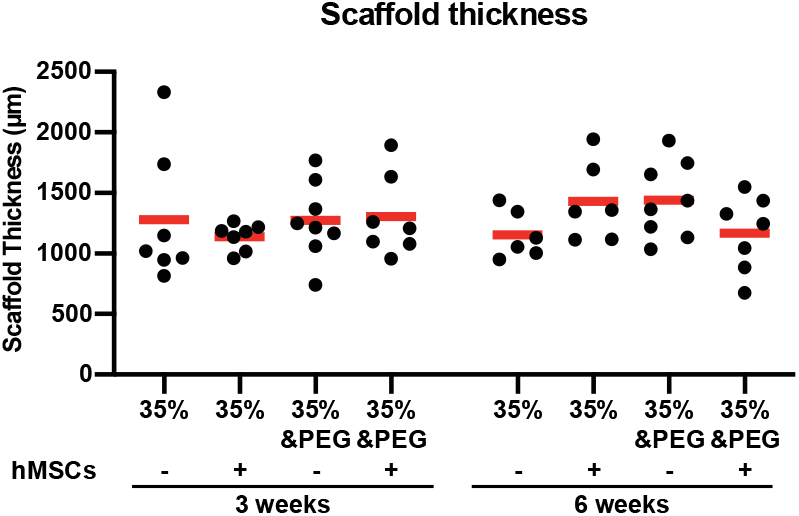
Scaffold thickness after explantation. Scaffold thickness was measured at the thickest section of a cryotome section taken from a random spot in the scaffold. n=6-8 for all conditions. One-way ANOVA, no statistical differences between groups.

**Supplementary movie 1. 3D view of nanoCT scan of a 20% scaffold**

**Supplementary movie 2. 3D view of nanoCT scan of a 20% &PEG scaffold**

**Supplementary movie 3. 3D view of nanoCT scan of a 35% scaffold**

**Supplementary movie 4. 3D view of nanoCT scan of a 35% &PEG scaffold**

